# STODE: A Deep Generative Framework for Continuous Spatiotemporal Dynamics in Spatial Transcriptomics

**DOI:** 10.1101/2025.11.24.690134

**Authors:** Koichiro Majima, Yasuhiro Kojima, Teppei Shimamura

## Abstract

**Motivation:** Spatial transcriptomics offers detailed molecular insights but struggles to capture continuous tissue dynamics due to experimental constraints and destructive sampling. Traditional optimal transport (OT) methods attempt to link time points but often rely on unimodal data and oversimplify complex developmental processes. We propose a spatiotemporal deep generative model that captures the continuous evolution of tissue, integrating both spatial and gene-expression dynamics. We first embed spatial transcriptomic particles into a latent space using a VAE and then learn time-dependent dynamics with a potential-guided neural ODE. This framework reveals diverse tissue behaviors, including growth patterns, spatial reorganization, and temporal gene-expression changes. A neural decoder estimates key behaviors like growth, disappearance, movement, and clustering, while a differentiable growth module models region appearance and disappearance.

**Results:** STODE is a deep generative model that reconstructs continuous spatiotemporal dynamics from discrete spatial transcriptomics measurements collected at multiple time points. On a complex synthetic dataset simulating gene-driven tissue morphogenesis and mouse midbrain development dataset, STODE significantly outperformed both linear interpolation and Entropic Optimal Transport models in predicting gene expression and spatial coordinates for a excluded timepoint. When applied to a mouse organogenesis dataset, the model’s backward simulations from late-stage embryos to early progenitors accurately reconstructed known developmental processes. The simulations correctly identified early pharyngeal arch derivatives as the populations occupying the largest area and captured the progressive differentiation of diverse cell types from multipotent neural crest and mesenchymal progenitors, demonstrating the model’s capability to reconstruct continuous and meaningful biological trajectories from discrete-timepoint spatial transcriptomes. In addition, the inferred factors reflected coordinated activation of morphogenetic programs, including cytoskeletal remodeling and cell-matrix interactions, and revealed transcriptional signatures associated with epithelial–mesenchymal transition and directed tissue migration, suggesting that the model effectively disentangled regulatory modules underlying the spatial organization and morphogenesis of craniofacial structures. The proposed model, STODE, will be broadly applicable to studies on embryogenesis, tissue regeneration, and disease progression, where reconstructing continuous spatiotemporal trajectories is essential for understanding biological mechanisms.

**Availability and Implementation:** The STODE model was implemented in Python using the PyTorch deep learning library. This code is available on GitHub at https://github.com/LzrRacer/STODE/.

## Introduction

During continuous biological processes such as tissue development and disease progression, cells differentiate into various distinct phenotypes during continuous biological processes such as tissue development and disease progression. Differentiation studies aim to understand cell state transitions in the cell state throughout this process, from the progenitor state to the post-differentiation state. This involves identifying the factors that regulate these changes and elucidating their underlying mechanisms of action.

The advent of spatial transcriptomics has enabled the measurement of gene expression within intact tissues along with spatial contexts, providing unprecedented insights into how molecular programs are organized across space (Ståhl *et al*., 2016). These technologies have advanced our understanding of tissue architecture, cellular interactions, and spatially patterned gene regulation. However, a fundamental limitation is their destructive nature; they only yield static snapshots of tissues at discrete time points. Thus, capturing the continuous dynamics of biological processes such as differentiation, migration, and tissue morphogenesis becomes inherently challenging as individual spots or cells cannot be tracked over time.

Although computational methods have been developed to infer temporal relationships from snapshot data, they often face significant challenges (Saelens *et al*., 2019). Trajectory inference methods may not adequately incorporate spatial information, and methods based on optimal transport (OT), which aim to align cell populations between time points, may struggle with the complexity of biological processes (Schiebinger *et al*., 2019). OT frameworks typically assume strict mass conservation and depend heavily on the choice of ground cost, making it difficult to model cell proliferation or disappearance, or the highly non-linear gene expression changes that drive development (Peyré *et al*., 2019).

To address these challenges, we developed STODE (Spatio Temporal latent dynamics with Ordinary Differential Equations), a potential-guided deep generative framework for reconstructing continuous cellular and tissue dynamics from discrete spatiotemporal transcriptome measurements. STODE combines a variational autoencoder (VAE) for non-linear dimensionality reduction with a neural ordinary differential equation (neural ODE) model that models temporal evolution in a unified latent–spatial state space. By first embedding each observational spot in spatial transcriptomics data an individual particle under a Lagrangian description and then modeling their trajectories using a neural ODE based on a gradient flow of the potential function, STODE captures the dynamic evolution of particle states. In this formulation, each particle resides within a continuous tissue field, carrying both the spatial position and molecular identity, and its motion represents the coordinated flow of gene expression and tissue morphology over time. This Lagrangian perspective enables STODE to model spatiotemporal dynamics in a broad range of time-series spatial transcriptomic datasets. This framework reconstructs diverse tissue behaviors, including growth patterns, spatial reorganization, and temporal changes in gene expression. Validation using a synthetic dataset and its application to a large-scale mouse embryogenesis dataset showed that STODE accurately interpolates unobserved states and performs backward simulations to uncover developmental processes, thereby linking transcriptional programs to dynamic tissue behaviors.

## Results

### A potential-guided VAE framework for spatiotemporal modeling

We developed a potential-guided deep learning framework to model and predict gene expression and tissue dynamics at unobserved timepoints, addressing the challenge in which experimental data are often discrete and sparsely sampled. As shown in the graphical model, our deep generative model learned the continuous dynamics of cell development from discrete spatiotemporal transcriptomic snapshots.

In this framework, based on the Lagrangian description, the continuum is explicitly treated as a collection of particles, and its overall behavior is characterized by tracking the trajectories of these particles. Each observational spot in spatial transcriptomics is regarded as an individual particle, representing an elementary unit of tissue that embodies both the spatial position and gene expression state, thereby serving as a local carrier of molecular and positional information within a continuous biological field. The model operates across two primary domains: the high-dimensional data space, representing the measured gene expression (**x**) and spatial coordinates (**s**), and a compressed, low-dimensional latent space (**z**) that captures the essential variations of expression profitless across spatiotemporal coordinates and evolution of these particles. Conceptually, this approach treats biological tissues as continuous systems in which local particles move and transform over time. Their collective trajectories capture smooth spatiotemporal flows of gene expression flow, enabling the reconstruction of coordinated molecular and morphological dynamics across diverse developmental and pathological contexts.

The core components of our model are a variational autoencoder (VAE), a potential field (*U*), and an ordinary differential equation (ODE) (Fig. S1) (Kingma *et al*., 2014; Chen *et al*., 2018). The VAE’s encoder maps high-dimensional gene expression data **x**_*t*_ onto the latent space representation **z**_*t*_. This is achieved through variational inference, in which the encoder network approximates the posterior distribution *p*(**z**|**x**) from the generative model with a tractable variational distribution *q*(**z**|**x**). The encoder outputs the mean (***µ***) and log variance (log ***σ***^2^) of a Gaussian distribution, from which the latent state **z**_*t*_ is sampled.

For each particle i at time t, we define a unified state vector **y**_*i,t*_ for each particle *i* at time *t* by concatenating its latent gene expression state **z**_*i,t*_ and its spatial coordinates **s**_*i,t*_:

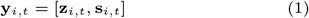

The temporal evolution of this field can be described using a Lagrangian formulation, following each particle as it moves through developmental time, rather than using the Eulerian view in which each particle is observed. This formulation allows us to track the continuous deformation and differentiation of tissues and capture the dynamic interplay between gene regulation and morphogenetic flow. The dynamics of this unified state are governed by a learnable, time-dependent potential function *U* that defines an energy landscape over the joint spatio-latent space, as follows:

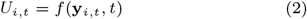

Here, the potential field acts as a guiding force for both the particle state and spatial coordinate. As **y**_*i,t*_ integrates gene expression and position, the potential simultaneously constrains how particles can change their transcriptional programs and move within the tissue space. The local minima of potential correspond to stable attractor states where particles with similar gene expression and positions accumulate, for example, differentiated cell types occupying specific tissue niches. Conversely, high-potential regions represent unstable or unlikely combinations of gene expression and spatial coordinates, discouraging biologically implausible transitions such as particles adopting an inconsistent phenotype or migrating to an incompatible spatial context. The temporal evolution of a particle’s state is then modeled as a continuous trajectory governed by an ODE, where the change in state over time is a function of the gradient of the potential function whose inputs are the state itself and the time point:

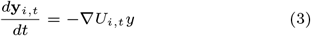

where the derivative *d***y**_*i,t*_*/dt* = −∇*U*_*i,t*_(*y, t*) describes the change in the unified state of particle i over time. In this formulation, the velocity of a particle in latent–spatial space is determined by its current state, the gradient of the potential, which acts as a force directing particles toward lower energy states; and explicit temporal effects. This enables the model to capture both transcriptional transitions and spatial rearrangements as smooth, continuous trajectories. Moreover, it facilitates inference of developmental processes, including changes in gene expression, spatial migration, and spatial expansion patterns, a inferred from vector field analysis (Fig. 1). This formulation allows the inference of developmental trajectories, including changes in gene expression, spatial migration, and population growth dynamics (Fig. 1).

**Fig. 1.**
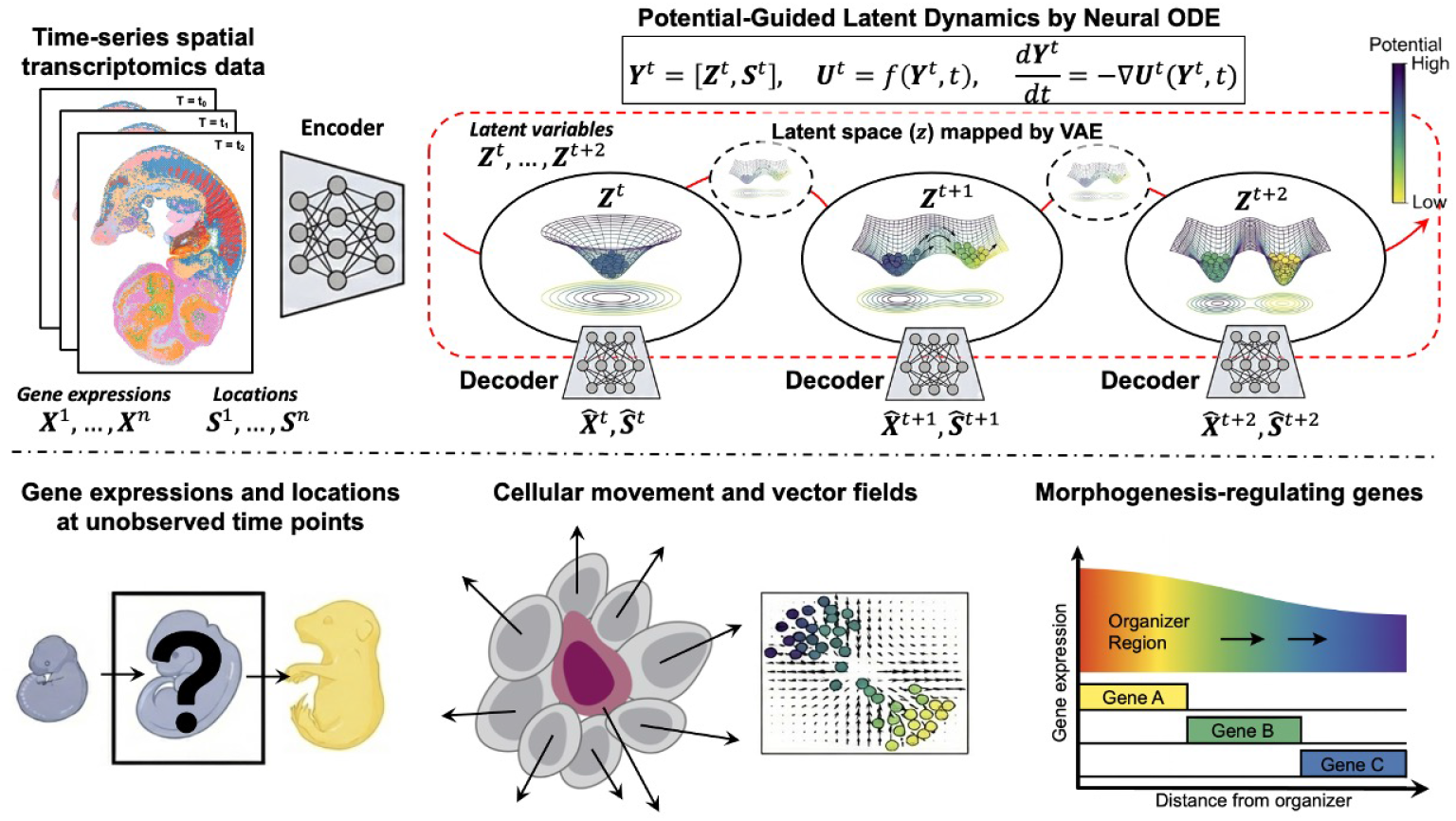
Conceptual overview of the STODE framework. **upper**, The model learns a time-varying potential energy landscape that guides the trajectory of particle states (spheres). A population of particles transitions from a single potential well (time *t*) to two distinct states (*t* + 1, *t* + 2) by following the gradients of the evolving landscape. **lower**, Key applications of the framework include: (Left) Predicting gene expression and morphology at unobserved time points; (Center) Inferring the velocity and migration trajectories of particle from the learned vector field; and (Right) Identifying genes whose expression correlates with morphological changes, ranking them as putative drivers of morphogenesis.

The parameters of the model’s dynamical components (e.g., potential function *U*) are jointly optimized by minimizing a composite loss function, as described in the following sections. This loss is designed to align the predicted dynamics with observed data snapshots while incorporating morphology-informed constraints that reflect the spatial organization of tissue. The total loss function is composed of several terms, each designed to capture different aspects of the model’s behavior. Each component is described in detail below:

#### Distribution Matching Loss

The primary objective of data fidelity was enforced using the sliced Wasserstein distance (*L*_SWD_) (Deshpande *et al*., 2018). This term quantifies the difference between the predicted and observed distributions over the joint particle state space defined by spatial coordinates and gene expression at each time point. Let **Y**_pred_ and **Y**_obs_ denote the sets of predicted and observed particle states, respectively. By projecting high-dimensional states onto a series of random 1D vectors (*θ*) and averaging the 1D Wasserstein-p distances (*W*_*p*_), this loss effectively quantifies the discrepancy in the joint state distribution between the predicted and observed particles without necessitating explicit matching of individual particles between corresponding time points.

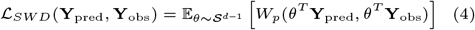

#### Temporal and Physical Regularization

To ensure that the learned dynamics are coherent, we introduce two regularization terms. The loss of temporal alignment, ℒ_align_, structures the latent space by training an auxiliary network, Regressor(·), to predict the biological time *t* from its latent code **z**_*t*_. This regularization encourages the latent space to reflect the flow of biological time, aligning changes in latent variables with temporal progression.

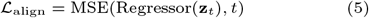

#### Force Consistency Regularization

The loss of force consistency ℒ_orce_ imposes a potential-guided constraint. This ensures that the velocity vector **v**_pred_ = −∇*U*_*t*_ predicted by the ODE network is consistent with the analytical force derived from the negative gradient of the potential field, **F**_analytic_ = −∇*U*_*t*_. It couples two networks and enforces a physically plausible dynamic.

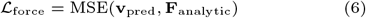

#### Total loss

The total loss ℒ_Total_ is the weighted sum of the four distinct terms:

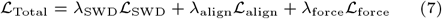

### STODE Accurately Predicts Tissue Dynamics in a Toy Model of Gene-controlled Morphogenesis

To validate the core capabilities of our framework, we first evaluated its ability to predict gene expression and spatial coordinates using a challenging synthetic dataset with a known ground truth. This toy dataset was designed to simulate anisotropic tissue growth, where upregulation of growth-accelerator genes strengthens the local velocity fields, leading to expansion along specified spatial axes, while repressor genes counteract this effect. This feedback between gene expression and tissue displacement is difficult to capture using simple interpolation models (see Methods). From a simulated five-point time course, the central timepoint (*t* = 3.0) was excluded from training, and the model was tasked with reconstructing the system state both gene expression and spatial coordinates using only the bracketing timepoints.

We compared the performance of STODE with that of two baselines: a simple linear interpolation model and a more advanced entropic optimal transport (OT) model that matches particle populations based on their gene expression profiles (Schiebinger *et al*., 2019). STODE was trained jointly on the bracketing timepoints, learning a latent representation and dynamic rules from a composite loss function that integrates latent dynamics, gene/spatial reconstruction, and KL-divergence terms.

Quantitative evaluation revealed that STODE achieved the lowest Mean Squared Error (MSE) for gene expression prediction and the lowest Root Mean Squared Error (RMSE) for spatial coordinate prediction (Table 1). Scatter plots of the predicted versus ground-truth spatial distributions showed a much tighter correlation compared with the baseline models (Fig. S2). Although targeted only a small set of genes associated with growth acceleration or repression, these results validate STODE’s ability to accurately capture gene expression transitions and complex spatiotemporal dynamics even using sparse, low-dimensional data.

**Table 1.** Quantitative comparison of interpolation accuracy on the toy dataset. Our STODE model demonstrates lower error in both gene expression and spatial coordinate prediction compared to baseline methods. Lower values are better.

| Metric | Linear | OT | STODE |
| --- | --- | --- | --- |
| Gene MSE ↓ | 0.85 | 0.62 | **0.21** |
| Spatial RMSE ↓ | 1.25 | 1.05 | **0.35** |

### Validation on Dorsal Midbrain Time-course Data

After confirming the ability of the model to accurately interpolate complex dynamics in a controlled setting, we applied it to a large-scale mouse midbrain development dataset to dissect its spatiotemporal dynamics. The assigned task was the same as in the previous section: an interpolation problem. That is, we inferred intermediate states using only E12.5 and E16.5 as inputs, whereas E14.5 was excluded from training.

Across all evaluation metrics, the VAE + latent ODE consistently outperformed the OT interpolation baseline, capturing the spatial organization patterns of the intermediate E14.5 stage without the tradeoffs observed in the moscot-based method (Fig. 2a, Fig. S4) (Klein *et al*., 2025).

**Fig. 2.**
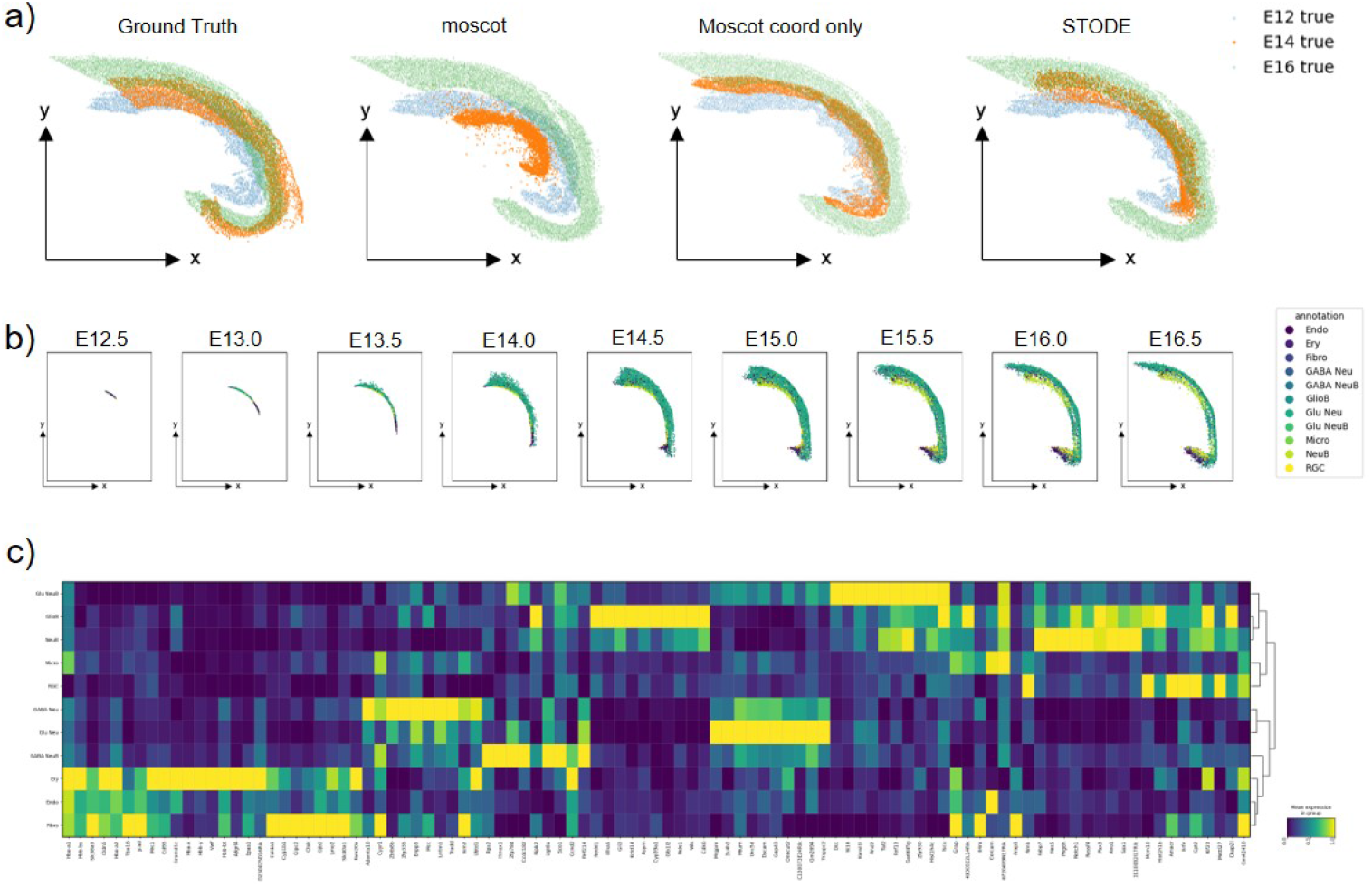
*In silico* Validation on Actual Dorsal Midbrain Time-course Data. **a**, Comparison of inferred developmental trajectories. From left to right: ground-truth reference, inference by MOSCOT using both the spatial coordinates and gene expression, MOSCOT using spatial coordinates only, and STODE (this study). Each panel shows the learned latent space projected onto the same 2D coordinate space in which the E16 data were observed (upper righit; color indicates biological time). **b**, Keyframes from the continuous reverse-time simulation showing morphological changes and cell type clustering from stages E11.5 to E6.5. **c**, Stacked bar chart showing the changing proportions of cell types over the simulated developmental timeline, capturing known organogenesis dynamics.

When using the standard moscot interpolation, the spatial accuracy was poor (OT coordinate MSE = 1.38; IoU = 0.059), reflecting distortions in coordinate predictions arising from the influence of gene expression on the joint cost. Adjusting moscot to emphasize spatial coordinates improved the geometric alignment (MSE = 0.413; IoU = 0.250) but substantially degraded expression accuracy (Poisson NLL = 7.80 *×* 10^6^). In contrast, our VAE + latent ODE achieved a low spatial error (MSE = 0.438) with the highest spatial overlap (IoU = 0.334) while simultaneously producing the best gene expression fit (MSE = 1.02 *×* 10^3^; Poisson NLL = 5.12 *×* 10^6^) (OT MSE measures the expected squared expression error under the OT coupling, and lower OT MSE or Poisson/NB NLL values indicate closer agreement between predicted and observed gene-expression profiles.) (Table S1).

These results demonstrate that latent ODE dynamics capture smooth temporal evolution in the joint expression-spatial manifold, avoiding the coordinate-expression tradeoff inherent in OT interpolation. By modeling continuous trajectories in the latent space, our method aligns more closely with developmental processes, producing more biologically coherent intermediate states.

Furthermore, we focused on cell types with increased proportions inferred from morphology (Fig. S3) and identified DEGs at E13. We found increased expression of *Hes5, Notch1* and *Febp7* in neuroblasts, and increased expression of *NDE1* in glioblasts (FIg. 2c) (Ohtsuka *et al*., 1999; Feng *et al*., 2000). In midbrain neuroblasts (Notch axis), *Notch1* maintains neural progenitors by activating the *Hes* family of genes and inhibiting premature neuronal differentiation(Shimojo *et al*., 2008).. *Hes5* acts as a Notch downstream effector, repressing proneural genes to sustain progenitor identity. Febp7 governs growth and differentiation signaling. In midbrain glioblasts, *NDE1* regulates mitotic spindle organization and neural progenitor division, both of which are essential for cortical development. In addition *NDE1* ensures the mechanical precision of microtubule–dynein–driven processes. This explains the increased proliferation of these cell types in these reagions. These findings suggest that STODE restores gene expression in a manner consistent with the experimental evidence.

### Application to Mouse Embryogenesis Data

We applied our spatiotemporal VAE–ODE (STODE) framework to the Mouse Organogenesis Spatiotemporal Transcriptomic Atlas (MOSTA) dataset (E9.5–E11.5), which includes tens of thousands of spatially resolved transcriptomes (Chen *et al*., 2022). After normalization, batch correction (ComBat), and selection of 2,000 informative genes, STODE embedded spatial transcriptome spots into a latent trajectory space constrained by neural dynamics.

The latent space resolved cell type–specific dynamics and reconstructed continuous transitions in the tissue architecture across destructive time-point samples. We visualized the spatial landscape of the latent space z and the potential, together with the cell-type classification (FIg. 3). For example, early trajectories were enriched for branchial arch structures, and the model captured the subsequent growth of the brain and surface ectoderm between E9.5 and E10.5, consistent with known embryological events such as somite formation and neuronal differentiation. Backward trajectories recapitulated conserved developmental principles: particles assigned to jaw and tooth primordia dominated the earliest simulated stages, reflecting derivation from the first pharyngeal arch; somitogenesis was captured by the emergence of dermatome-like populations at intermediate stages; and the populations of particles assigned to neural crest stem cells and mesenchymal cells increased as trajectories converged toward progenitors, consistent with their broad differentiation potential whereas brain and surface ectoderm contributions increased at later stages. Gene-level predictions were biologically coherent; Fez1 was upregulated in neural tissues during axonal development, whereas hepatic regions showed the induction of PCSK9, consistent with early metabolic programming (Fig. S5).

**Fig. 3.**
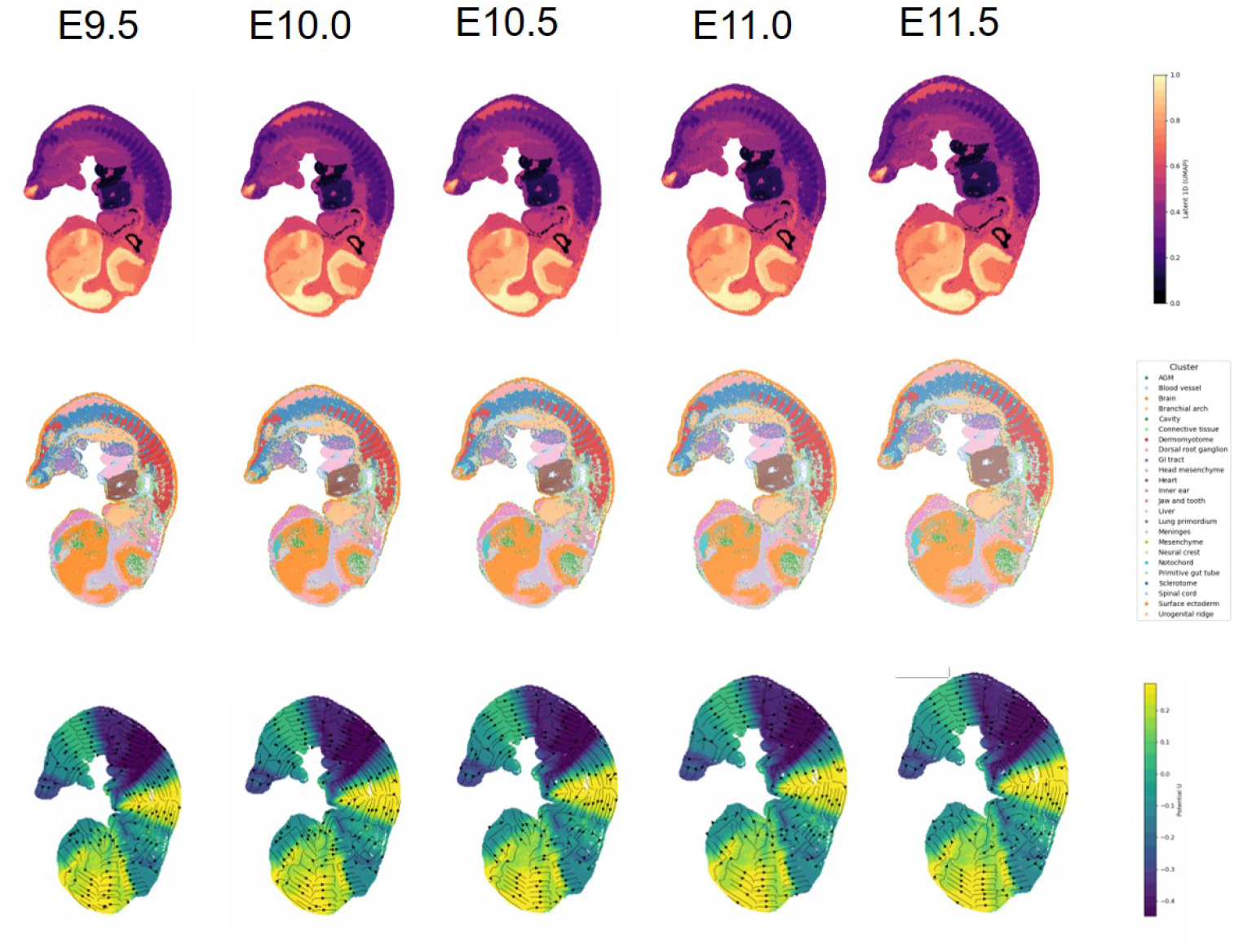
*In silico* simulation of mouse organogenesis by STODE. Keyframes from the continuous reverse-time simulation showing morphological changes **upper**, Latent variable z learned by VAE. **middle**, Cell type clustering in unobserved time points. **lower**, Streamline plot of the potential calculated by the model and the direction of movement of each particle obtained by differentiation. The color scale represents the divergence of the inferred vector field, where positive values indicate expansion and negative values indicate contraction.

Overall, STODE integrated spatial and transcriptomic information to reconstruct gene regulation, and lineage hierarchies. Notably, retrograde simulations inferred intermediate states that were absent from experimental sampling, providing mechanistic hypotheses for unobserved but biologically plausible developmental transitions.

### Analyzing Spatial Growth via Vector Field Divergence

Our trained generative model provides not only a means of simulating developmental trajectories in gene expression space but also a framework for analyzing the underlying dynamics of tissue morphogenesis. The model learns a continuous spatiotemporal vector field representing the particle velocities in both spatial and latent spaces.

The state of any particle at a biological time *t* is defined by a complete state vector **Y**(*t*) = [**s**(*t*), **z**(*t*)], where 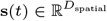 are the spatial coordinates and 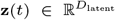 is the latent vector representing the transcriptional state of the particle (derived from the gene expression matrix, conventionally denoted as *X*). The trained ODE model *f*_ODE_ predicts the instantaneous velocity of the state vector

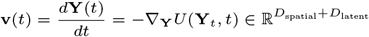

To specifically analyze the physical organization of tissue, we can isolate the spatial component of this velocity field,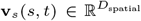, which describes the movement of particles in physical space. This extraction is performed through a projection matrix 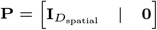.

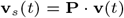

The divergence of the spatial velocity field div(**v**_*s*_) = ∇_*s*_ · **v**_*s*_ quantifies the local tissue expansion and compression. Positive divergence (∇ · **v**_*s*_ *>* 0) identifies the sources of particle flow, corresponding to regions of net volumetric growth or dispersion. Negative divergence (∇ · **v**_*s*_ *<* 0) marks sinks, indicating local particle condensation. By computing the divergence across space and time, we generated dynamic maps of morphogenetic sources and sinks, offering quantitative insight into the principles of tissue development.

By computing the divergence across the spatial domain at various time points, we generated dynamic maps that visualized how these morphogenetic source and sink regions evolved, offering key insights into the principles governing tissue development (Fig. 3c).

We computed the mean divergence of major annotated tissues across the continuous developmental timeline inferred by the model (Fig. 4). Our analysis revealed consistently high divergence values in the developing brain and heart throughout early organogenesis (E7.5-E11.5), quantitatively capturing the rapid expansion characteristic of these organs (Xie *et al*., 2025; Ermakova *et al*., 2018). The branchial arch and liver also showed significant, though slightly lower, divergence, consistent with their roles as major sites of growth and morphological change during this period (Depew *et al*., 2005; Lee *et al*., 2012). In contrast, tissues such as the notochord, sclerotome, and connective tissue exhibited markedly lower divergence, suggesting more stable tissue flux and less volumetric expansion (Bellomo *et al*., 1996). These results demonstrate the ability of our model to visualizes and quantify fundamental morphogenetic events, providing a dynamic signature of tissue growth and reorganization that aligns with established developmental biology.

**Fig. 4.**
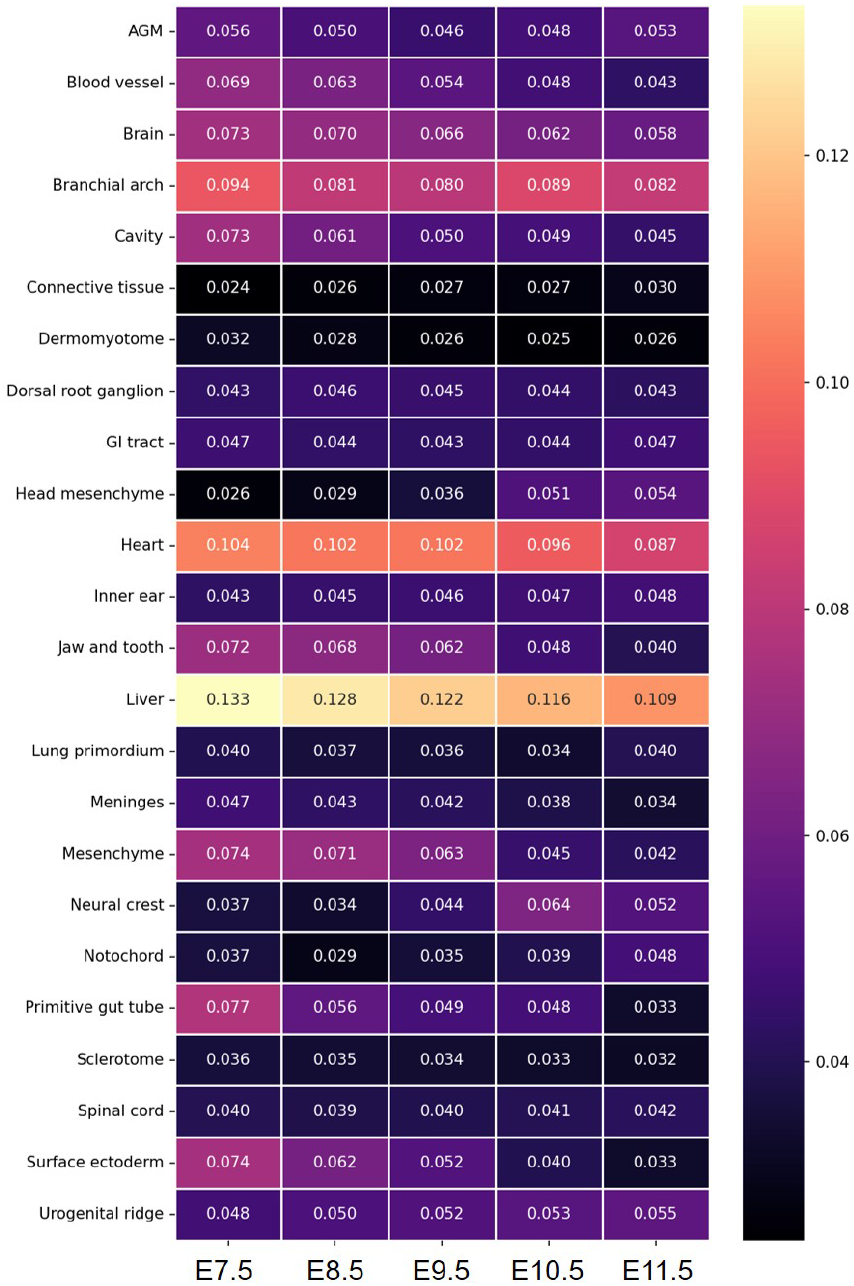
Quantitative analysis of tissue dynamics via divergence. Heatmap of the mean absolute divergence of the learned velocity field, calculated for major cell types across the simulated timeline (E7.5-E11.5).

### Identifying Genetic Drivers of Tissue Morphogenesis

To connect the emergent physical dynamics of tissue formation with their underlying molecular regulators, we developed an approach to identify genes whose expression patterns are associated with learned tissue velocities. For each particle, we computed the magnitude of its velocity vector ∥**v**∥ as predicted by the learned dynamics field. We then systematically calculated the correlation between the velocity magnitude and the expression level of each gene across all particles at each time step. This analysis allowed us to identify genes whose expression was significantly correlated with particle movement, thereby identifying them as potential drivers of morphogenesis.

Our analysis identified a cohort of genes whose expression levels strongly correlated with high-velocity tissue movement (Fig. 5). Notably, this list includes genes encoding essential cytoskeletal proteins such as cardiac actin (*Actc1*) and smooth muscle actin (*Acta2*), which form the core machinery for cell contractility and motility (Matsson *et al*., 2008; Rockey *et al*., 2013; Brooks *et al*., 2010). Furthermore, we identified genes involved in the regulation of the cytoskeleton, such as *Nhs*, which plays a role in actin remodeling. Importantly, our analysis also highlighted key developmental transcription factors, including *Prrx1*, which is a critical regulator of cell migration in mesenchymal tissues (Du *et al*., 2021; Feldmann *et al*., 2021). The identification of these biologically plausible candidates validates our approach and demonstrates the capacity of the model to forge a direct link between macroscopic tissue dynamics and their specific molecular drivers, offering a powerful tool for generating new, testable hypotheses regarding the genetic control of development.

**Fig. 5.**
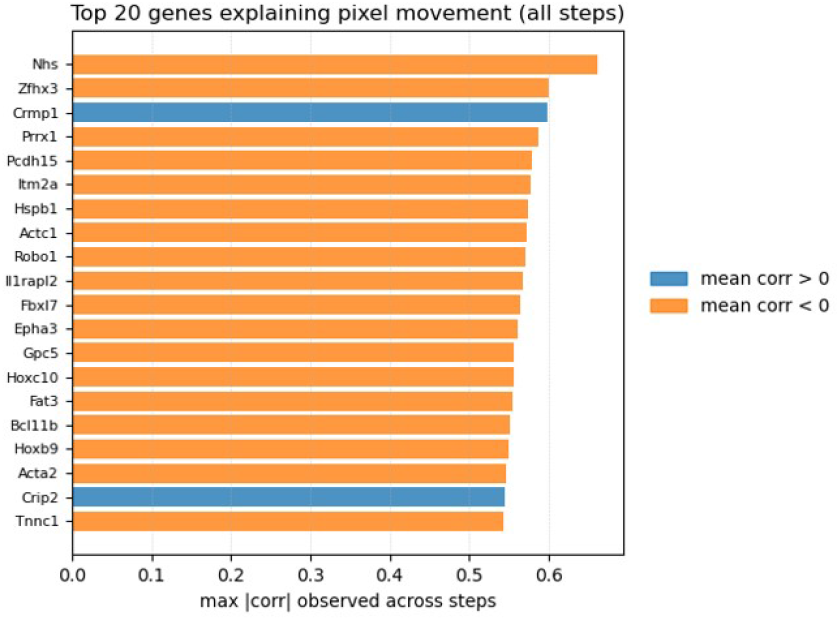
Genes correlated with inferred particle movement. The top 20 genes with the highest correlation between their expression and the inferred particle displacement vectors.

## Discussion

In this study, we introduce STODE, a spatiotemporal VAE–ODE framework that integrates transcriptomic and spatial measurements into a unified latent representation governed by continuous dynamics. Conceptually, STODE adopts a Lagrangian description of tissue development, in which each observational spot is treated as a material particle whose trajectory encapsulates the local molecular and positional evolution. This particle-based view enables the reconstruction of coherent developmental flows using static and destructive timepoint measurements.

By leveraging a potential-based formulation and potential-guided regularization, STODE reconstructs biologically meaningful developmental trajectories, while preserving spatial and temporal consistency. This approach directly addresses a central limitation of spatial transcriptomics: although technologies such as Stereo-seq provide high-resolution spatial snapshots, they cannot directly capture continuous processes such as particle migration, proliferation, or morphogenesis (Chen *et al*., 2022). Our framework bridges this gap by inferring smooth spatiotemporal trajectories that approximate the underlying dynamic fields of tissue development.

Compared with interpolation or OT-based approaches, STODE offers several advantages. First, it jointly embeds gene expression and spatial coordinates into a low-dimensional latent space, ensuring that the temporal dynamics reflect both molecular and morphological changes. Second, the incorporation of a potential field and an ODE solver enables the model to simulate forward and backward trajectories, filling in unobserved intermediate states. This retrograde inference capability provides mechanistic hypotheses for unobserved particle states, which cannot be achieved using standard OT formulations. Finally, the framework naturally yields interpretable outputs such as vector field divergence and gene–velocity correlations that directly link molecular drivers to tissue-scale morphogenesis.

Applications to both synthetic benchmarks and the MOSTA dataset demonstrated that STODE can reconstruct known embryological events such as somite formation, pharyngeal arch development, and brain growth. Importantly, the model also generated novel, testable predictions such as those for specific transcription factors and cytoskeletal genes correlated with particle displacement. These results suggest that STODE can serve as a general computational paradigm for bridging the gap between transcriptional regulation and morphogenetic outcomes.

## Conclusion

In summary, we present STODE, a deep generative model for reconstructing continuous tissue development from discrete spatial transcriptomic data. Our framework integrates a variational autoencoder with neural ODE dynamics guided by a potential energy landscape, thereby enabling the inference of both forward and backward developmental trajectories. Across synthetic and biological datasets, STODE outperformed existing interpolation and OT-based methods in reconstructing gene expression and spatial dynamics. Beyond predictive accuracy, the model provides interpretable mechanistic insights, linking gene expression programs to emergent physical processes, such as tissue expansion, migration, and lineage specification. We anticipate that STODE will be broadly applicable to studies on embryogenesis, tissue regeneration, and disease progression, where reconstructing continuous spatiotemporal trajectories is essential for understanding biological mechanisms.

## Limitations of the Study

While STODE provides a powerful framework for modeling spatiotemporal dynamics, several limitations should be noted. First, like most neural generative models, it requires careful hyperparameter tuning and substantial computational resources, which may limit accessibility for very large datasets. Second, the reliance on destructive sampling means that the model infers rather than directly observes temporal continuity, and errors in data preprocessing (e.g., batch correction or gene selection) may propagate into trajectory predictions. Third, although retrograde inference offers biologically plausible reconstructions, these remain computational hypotheses that require independent experimental validation, such as lineage tracing or live imaging. Finally, the current framework assumes smooth dynamics, which may underrepresent rare, abrupt developmental transitions or stochastic events. Future work will focus on extending the model to integrate multi-modal data (e.g., epigenomics, proteomics) and to incorporate uncertainty quantification, enabling more robust and experimentally grounded inference of complex tissue dynamics.

## Methods

### Generative Model for Sequential Latent State Transition

This section describes the generative model for time-series spatial transcriptomes. This probabilistic model uses latent variables, *z*_*t*_ ∈ *R*^*m*^, where *m* is the dimension of the latent particle state space and *t* is the time of the particle state.

Each lineage (a population of particles integrated with the same DNA barcode) begins with a unique progenitor particle state that diffuses within the latent space over time. This expression was derived from the latent state after repeated diffusion of the particle state up to the observation time. The latent particle state is represented by ***z*** and follows a Gaussian prior distribution. The dynamics ***z***_*t*_ − ***z***_*t*−1_ represents the transition of the particle state during a certain period from time point ***t*** and is expressed as the difference in the particle state between specific time points. These values followed a Gaussian prior distribution:

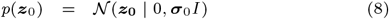

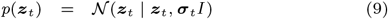

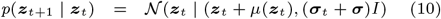

### Data Preprocessing and Batch Correction

A publicly available mouse organogenesis spatiotemporal transcriptomics dataset was used for all analyses. The raw data were loaded as AnnData objects, and a comprehensive preprocessing pipeline was applied using the Scanpy v1.9 package (Wolf *et al*., 2018).

The initial quality control involved filtering out particles expressing fewer than 100 genes, and genes detected in fewer than 10 particles. To mitigate the technical artifacts arising from measuring particles at different developmental stages, we treated each timepoint as a distinct batch. We applied the ComBat algorithm (`sc.pp.combat`) to the log-normalized expression values to correct for batch effects (Johnson *et al*., 2007). This procedure yielded a continuous batch-corrected expression matrix suitable for downstream modeling with a Gaussian likelihood. Finally, we selected a common set of 2,000 highly variable genes (HVGs) across all timepoints for subsequent modeling and analysis. The resulting preprocessed AnnData object was saved for model training.

### A Spatiotemporal Generative Model for Particle Dynamics

To model the continuous dynamics of embryonic development, a deep generative framework was developed that learns the joint distribution of gene expression and spatial coordinates over time. Our formulation is inspired by the **Lagrangian description** of continuum mechanics, in which a system is represented as an ensemble of particles whose individual trajectories collectively define the evolution of the whole system. By this analogy, each observational spot in spatial transcriptomics is regarded as a particle carrying molecular and positional information within a continuous biological field.

Conceptually, our model is similar to frameworks such as PI-SDE and scNODE, which model particle dynamics in a learned low-dimensional space (Jiang *et al*., 2024; Zhang *et al*., 2024). Specifically, our model integrates a VAE for dimensionality reduction with a system of neural ODEs that govern the dynamics in the learned latent space (Fig. 1).

#### Variational Autoencoder for particle State Representation

We employed the VAE to learn a compressed, low-dimensional representation of the particle state from high-dimensional gene expression data. The VAE comprises two multi-layer perceptron (MLP) networks:

1. An **encoder**, *q*_*ϕ*_(**z**|**x**), which maps the input gene expression **x** ∈ ℝ^*p*^ to the parameters of a diagonal Gaussian distribution in the latent space, 𝒩 (*µ, σ*^2^), where **z** ∈ ℝ^*d*^ is the latent representation.
2. A **decoder**, *p*_*θ*_(**x**|**z**), which reconstructs the original gene expression profile from a point in the latent space.

Our specific architecture utilizes an encoder and a decoder with hidden layers of size (128, 64) mapped to a latent space of dimension *d* = 8. The model is optimized by minimizing the composite loss function *L*_VAE_, which consists of a reconstruction term and a Kullback–Leibler (KL) divergence term to regularize the latent space:

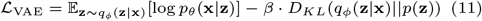

Given our continuous batch-corrected input data, we used a Gaussian likelihood for the reconstruction term, which is equivalent to minimizing the mean squared error (MSE). The term *p*(**z**) is a standard Gaussian prior 𝒩 (0, *I*), and *β* is a weight hyperparameter for the KL term.

#### Neural ODE for particle Dynamics

We modeled the developmental process as a continuous trajectory in a joint state space defined by particle spatial coordinates **s** ∈ ℝ^2^ and its latent representation **z** ∈ ℝ^8^. The evolution of this joint state was governed by a system of ODEs. The central hypothesis inspired by potential-guided models is that the velocity of a particle is driven by the negative gradient of a learned time-varying potential energy function Φ(**s, z**, *t*). This potential function defines the underlying energy landscape of the particle development. The system is defined as follows:

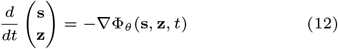

This system comprises two key neural network components.

- A **SpatioTemporal Potential network**, Φ_*θ*_(**s, z**, *t*), which learns a scalar potential energy landscape that varies with space, latent state, and time.
- A **Time-Aware ODE network**, *f*_*ω*_, which approximates the velocity vector field, modeling a form of damped dynamics.

### Model Training and Optimization

The model is trained using a two-stage process. First, the VAE is pre-trained on the entire preprocessed dataset for 100 epochs to establish a stable latent space. Following pre-training, the VAE weights are frozen, and the dynamical components (Φ_*θ*_ and *f*_*ω*_) are trained end-to-end for 50 epochs using the AdamW optimizer with a learning rate of 5 *×* 10^−4^. The composite loss function for the second stage, ℒ_Total_, is designed to ensure accurate distribution matching and physically plausible dynamics.

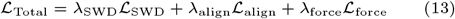

The loss components are defined as follows.

- ℒ_**SWD**_: The sliced Wasserstein distance (SWD) between the distribution of particle states predicted by the model at time *t*_*k*−1_ and the observed data distribution at *t*_*k*−1_. This aligns the marginal distributions of the learned process with the data.
- ℒ_**align**_: A time-alignment loss that enforces temporal coherence in the latent space by training an auxiliary MLP to regress biological time from the latent state **z**.
- ℒ_**force**_: A force-consistency loss that regularizes the ODE network by encouraging its predicted velocity field *f*_*ω*_ to align with the negative gradient of the learned potential, −∇Φ_*θ*_. This is conceptually similar to potential-guided approaches that enforce consistency with underlying principles. To stabilize the training further, an additional convergence term can be incorporated.

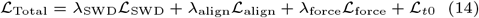
- ℒ_*t*0_: A convergence loss that penalizes variance and deviation from a specified target state for all trajectories at the initial timepoint *t*_0_. The target state at *t*_0_ is determined dynamically by identifying the latent coordinates of the center of mass or the most undifferentiated particles (putative progenitors) at the latest observed timepoint, using differentiation-degree inference algorithms such as *CytoTrace* (Gulati *et al*., 2020).

### Synthetic Data Generation (Toy Experiment)

To evaluate STODE under controlled and interpretable conditions, we constructed a synthetic dataset that simulated anisotropic tissue growth modulated by gene expression feedback. The simulation represents a two-dimensional tissue comprising *N* particles, each characterized by a spatial coordinate **s**_*i*_(*t*) ∈ ℝ^2^ and a gene expression vector **x**_*i*_(*t*) ∈ ℝ^*G*^, where *G* is the number of genes. In this setup, all genes were categorized into three functional types: growth accelerators, growth repressors, and neutral reference genes.

#### Spatial Dynamics

Particle positions evolved according to the time-dependent anisotropic growth field

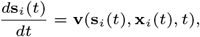

where **v**(·) is the growth velocity function influenced by local gene expression. For each simulation step, the velocity was updated as follows:

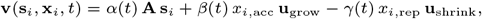

where **A** encodes baseline isotropic growth, **u**_grow_ and **u**_shrink_ are directional unit vectors, and *x*_*i*,acc_ and *x*_*i*,rep_ denote expression levels of the accelerator and repressor genes, respectively. The coefficients *α*(*t*), *β*(*t*), and *γ*(*t*) modulate the global influences of the baseline, accelerating, and repressing components over time, respectively.

#### Gene Regulatory Feedback

Gene expression is dynamically updated as a function of spatial context and developmental time. For simplicity, we define

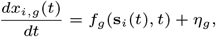

where *f*_*g*_ represents gene-specific regulatory functions and 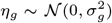 introduces stochastic transcriptional noise. In this toy model, accelerator genes are upregulated in the frontier regions (particles with high projected barycentric coordinate values), whereas repressor genes are enriched in the posterior regions:

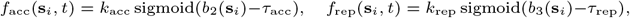

where *b*_2_(**s**_*i*_) and *b*_3_(**s**_*i*_) are the barycentric coordinates corresponding to the two growth fronts.

#### Numerical Simulation

The coupled system of ordinary differential equations for **s**_*i*_(*t*) and **x**_*i*_(*t*) is integrated using an explicit Runge–Kutta (RK4) scheme over *T* discrete time steps. At each observation time *t*_*j*_ ∈ {0, 1, 2}, the particle positions and the corresponding gene expression states were simulated.

### Simulation and Downstream Analysis

#### Backward Simulation and particle Merging

To reconstruct the developmental history, we performed backward simulations starting from the observed particle state at the latest timepoint (e.g., E11.5). Particle state trajectories were generated by numerically integrating the learned ODE system backward in time using a custom Euler integrator. Our simulation incorporates a dynamic hlparticle merging mechanism to account for the natural increase in the particle density at earlier developmental stages. During integration, the particles that fall within the same bin of a predefined spatial grid are merged into a single particle by averaging their state vectors.

#### Temporal Clustering and Annotation Mapping

To analyze the emergent particle populations within the simulated trajectories, we sampled the particles at discrete time points and performed clustering. We used two complementary approaches: (1) *de novo* clustering using the k-means algorithm on the decoded gene expression data and (2) a novel annotation strategy that maps simulated particles to known biological cell types by finding the nearest observed cell-type centroid in the VAE’s latent space. This allows for a direct biological interpretation of simulated particle populations and their temporal evolution.

#### Annotation-based Cluster Assignment

To assign biological identities to the simulated particles, we mapped them to the nearest annotated cell type from the observed data within the VAE’s latent space.

First, for each ground-truth annotation *a*_*k*_ present in the set of all observed annotations *A*, we computed a centroid vector **c**_*k*_ in the *D*_*z*_-dimensional latent space. This centroid represents the average latent representation of all observed particles belonging to the annotation. The calculation was performed using the VAE’s encoder *E*_*ϕ*_:

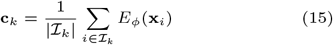

where ℐ_*k*_ = {*i*|annotation(**x**_*i*_) = *a*_*k*_} denotes the index set of the observed particles with annotation *a*_*k*_.

Next, for each simulated particle with a decoded expression profile 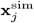, we obtained the latent representation 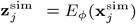. The cluster label *L*_*j*_ for this simulated particle was then assigned by identifying the annotation centroid closest to its latent representation, which is measured using the Euclidean distance

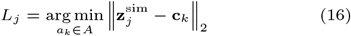

This process effectively partitioned the simulated particles into groups that correspond to the known biological cell types observed in the experimental data, thereby enabling a direct comparison and analysis of their temporal dynamics.

#### Latent Space and Gene-Displacement Analysis

To gain insights into the learned developmental manifold, we characterized the temporal evolution of each cluster within the latent space by tracking the mean and variance of their latent coordinates over time. Furthermore, to link gene expression to particle dynamics, we computed the Pearson correlation between gene expression at time *t*_*i*_ and the magnitude of its simulated spatial displacement between *t*_*i*_ and *t*_*i*+1_. This analysis identified genes whose expression levels were predictive of particle motility.

## Code Availability

The code used for all the analyses and simulations is available at [Repository URL].

## Resource availability

### Lead contact

Further information and resource requests should be directed to and will be fulfilled by the lead contact, Teppei Shimamura.

## Materials availability

This study did not generate new unique reagents.

## Data and code availability

The mouse stereo-seq atlas data were retrieved from Chen *et al*. and are available at https://db.cngb.org/stomics/mosta/ (Chen *et al*., 2022). The Axolotl stereo-seq atlas data were retrieved from Wei *et al*. and are available at https://db.cngb.org/stomics/artista/ (Wei *et al*., 2022). The STODE model was implemented in Python using the PyTorch deep-learning library, and the code is available at https://github.com/LzrRacer/STODE/ (Paszke *et al*., 2019).

Any additional information required to reanalyze the data reported in this paper is available from the lead contact upon request.

## Author contributions statement

K. Minoura conceived the idea for this study. K. Majima designed and performed the experiments under the supervision of Y.K. and T.S. All authors have read and approved the final manuscript.

## Acknowledgments

This work was supported by multiple sources. Grant-in-Aid for Transformative Research Areas (Platforms for Advanced Technologies and Research Resources) (grant no. 22H04925) and Grant-in-Aid for Transformative Research Areas (A) (grant no. 23H04938) were provided by the Japan Society for the Promotion of Science (JSPS). Additional support was received from Brain/MINDS Health and Diseases (grant no. JP25wm0625519), the Interdisciplinary Cutting-edge Research (grant no. JP25wm0325068), the Moonshot R&D program (grant no. JP25zf0127012), and the Advanced Genome Research and Bioinformatics Study to Facilitate Medical Innovation (GRIFIN) (grant no. JP25tm0424226) from the Japan Agency for Medical Research and Development (AMED). The Moonshot R&D program (grant no. JPMJMS2025) also contributed through the Japan Science and Technology Agency (JST). Further support came from the Medical Research Center Initiative for High Depth Omics and Multilayered Stress Diseases at the Institute of Science Tokyo. Supercomputing resources were provided by the Shirokane supercomputer at the Human Genome Center of the University of Tokyo and the TSUBAME3.0 supercomputer at the Institute of Science Tokyo.

---

**Table S1.** Quantitative comparison of interpolation accuracy on the embryonic midbrain dataset. Quantitative comparison of interpolation accuracy on the embryonic midbrain dataset. The table lists coordinate and gene expression prediction errors, as well as spatial overlap, for three interpolation strategies: standard moscot OT, moscot OT with coordinate-only emphasis, and STODE. Metrics include OT coordinate MSE, coupling-weighted expression MSE (log1p), Poisson NLL, and coordinate occupancy IoU.

| Model | OT coord. MSE ↓ | Expr. MSE (log1p) ↓ | Poisson NLL ↓ | Coord. IoU ↑ |
| --- | --- | --- | --- | --- |
| moscot_OT | 1.379 | 1267.96 | $7.95 \times 10^6$ | 0.059 |
| moscot_OT_coord_only | 0.413 | 1251.73 | $7.80 \times 10^6$ | 0.250 |
| **STODE** | **0.438** | **1021.33** | $5.12 \times 10^6$ | **0.334** |

**Fig. S1.**
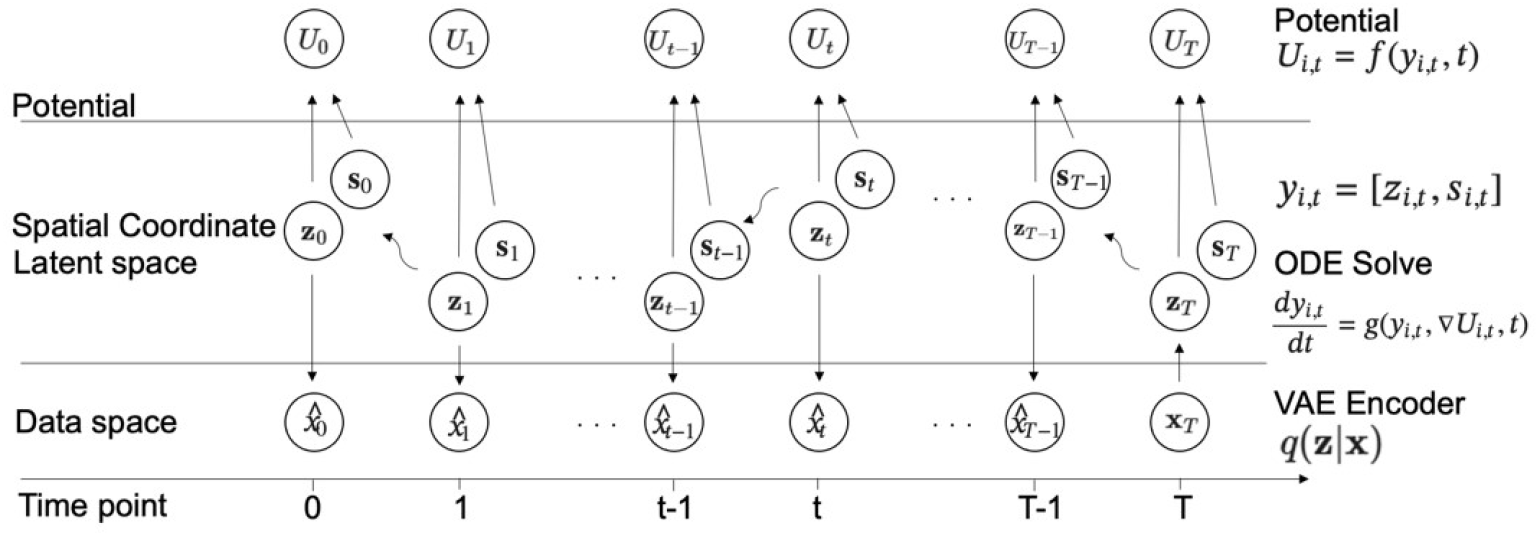
Graphical model of the spatiotemporal framework. The model integrates a Variational Autoencoder (VAE) with a potential-guided Neural Ordinary Differential Equation (NODE). High-dimensional gene expression data (*x*) is encoded into a latent space (*z*). The latent state is combined with spatial coordinates (*s*) to form a unified state vector, *y* = [*z, s*]. The continuous dynamics of this state are governed by a learnable potential (*U*) and a NODE, *dy/dt* = −∇*U*, which models the velocity of the particle state.

**Fig. S2.**
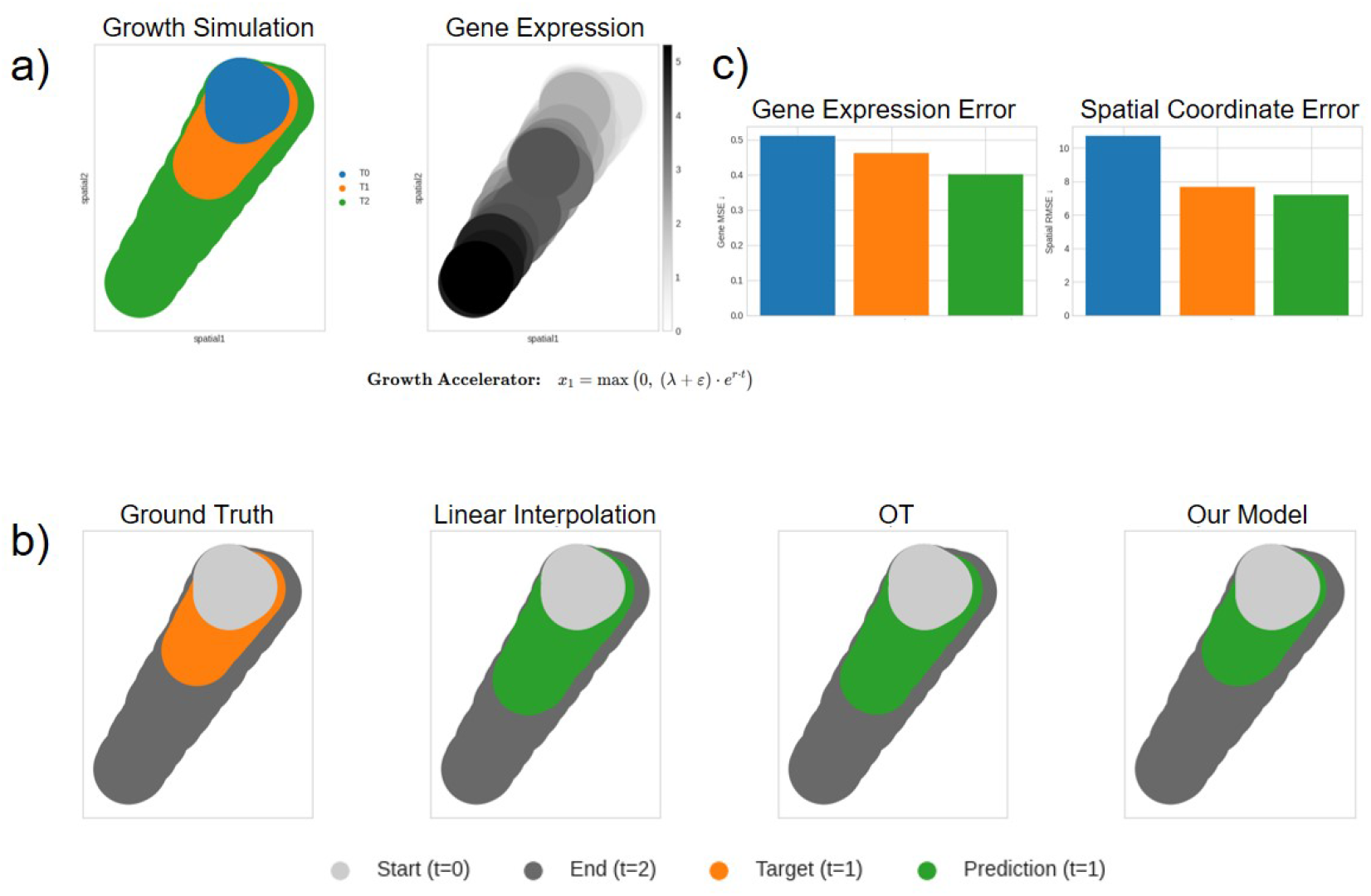
Performance on a simulated non-linear growth dataset. **a**, The toy dataset simulates non-linear tissue growth driven by a “Growth Accelerator” gene with exponential expression over time. **b**, Visual comparison of model interpolation at t=1. The prediction of Linear Interpolation or Entropic Optimal Transport (OT) and the proposed model (green), ground truth (orange). **c**, Quantitative error for the interpolation task.

**Fig. S3.**
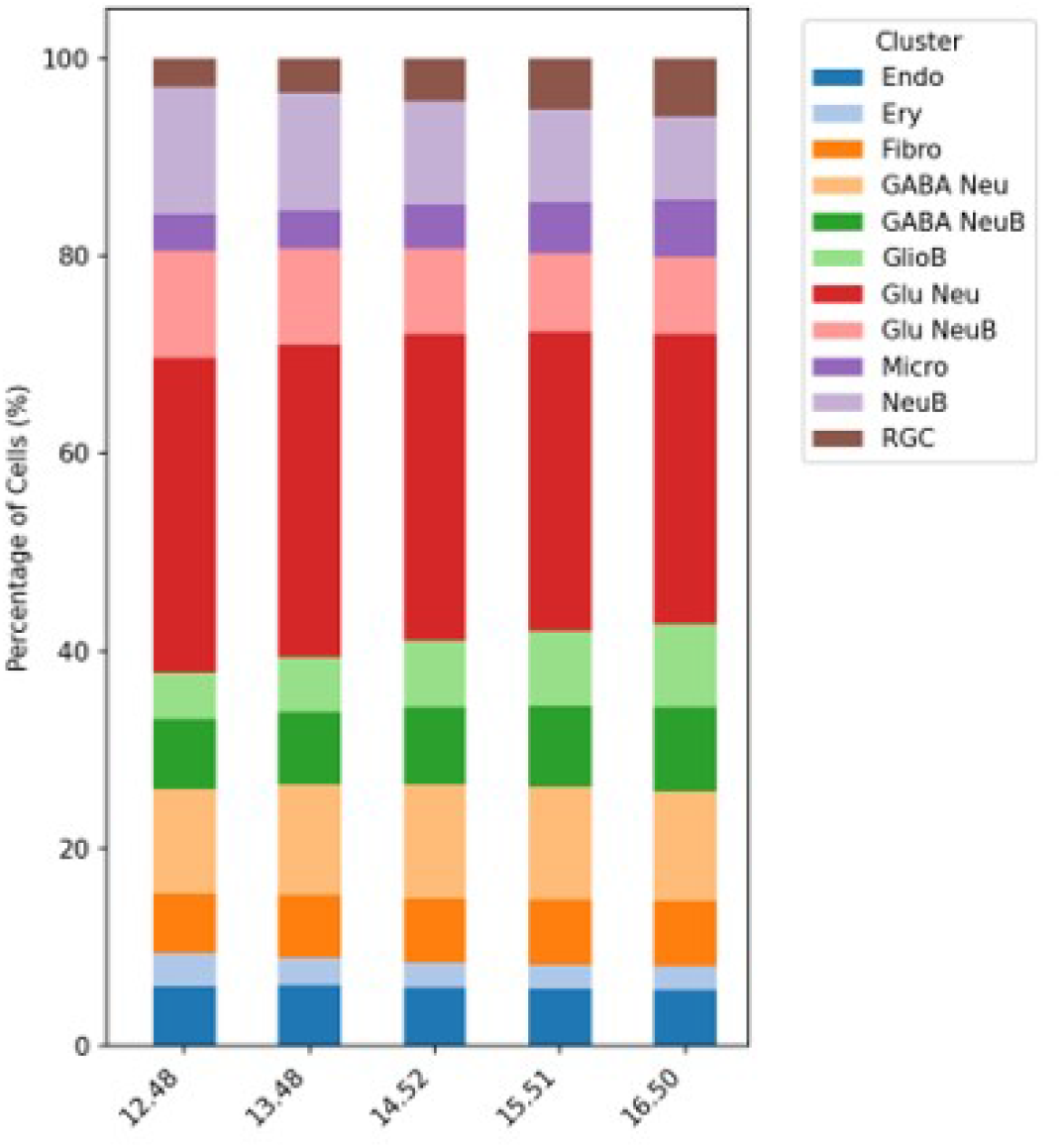
Proportion of each cell type in the whole tissue. Cell types are assigned based on the inferred latent variables, and the proportion of pixels of that cell type in the whole tissue is displayed in a bar chart.

**Fig. S4.**
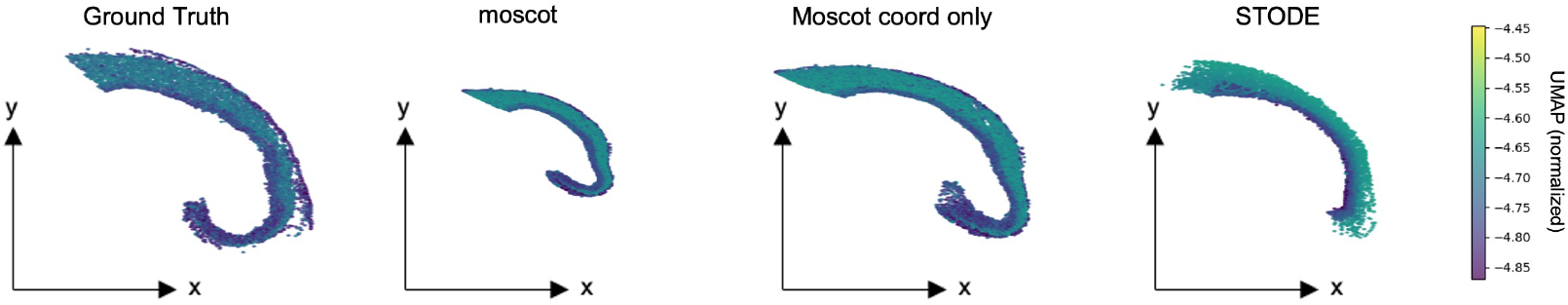
Comparison of inferred developmental trajectories at the intermediate time point (E14.5). From left to right: ground-truth observation, inference by MOSCOT using both spatial coordinates and gene expression, MOSCOT using spatial coordinates only, and STODE (this study). Each panel shows the reconstructed spatial configuration in the same two-dimensional coordinate system as the observed data, with cells colored by the UMAP_1_ values (normalized across methods) to highlight differences in the recovered manifold structure.

**Fig. S5.**
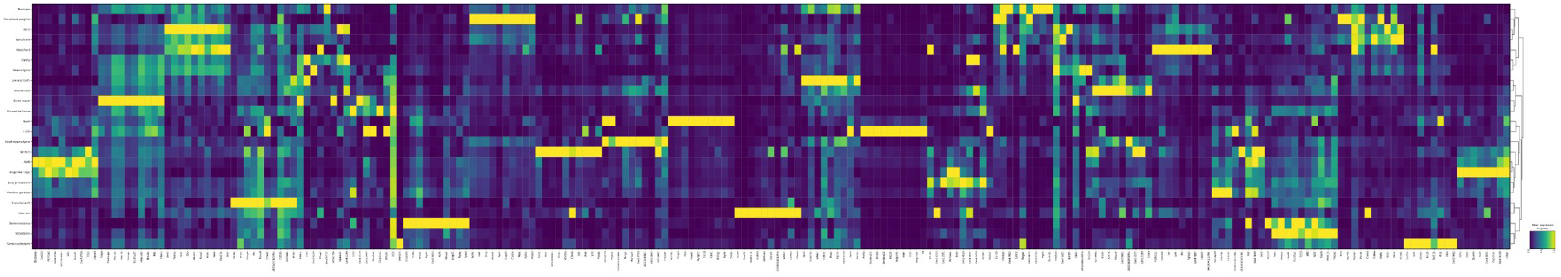
Predicted marker gene expression validates simulated cell types. Heatmap of predicted expression for known marker genes across simulated cell types at embryonic day 6.5 (E6.5).

